# HDAC3 regulates senescence and lineage specificity in non-small cell lung cancer

**DOI:** 10.1101/2020.10.14.338590

**Authors:** Lillian J. Eichner, Stephanie D. Curtis, Sonja N. Brun, Joshua T. Baumgart, Debbie S. Ross, Tammy J. Rymoff, Reuben J. Shaw

## Abstract

Transcriptional deregulation is a common feature of many cancers, which is often accompanied by changes in epigenetic controls. These findings have led to the development of therapeutic agents aimed at broad modulation and reprogramming of transcription in a variety of cancers. Histone Deacetylase 3, HDAC3, is one of the main targets of HDAC inhibitors currently in clinical development as cancer therapies, yet the *in vivo* role of HDAC3 in solid tumors is unknown. Here, we define the role of HDAC3 in two genetic engineered models of the most common subtypes of Kras-driven Non-Small Cell Lung Cancer (NSCLC), Kras^G12D^, STK11^−/−^ (KL) and Kras^G12D^, p53^−/−^ (KP), where we found that HDAC3 is strongly required for tumor growth of both genotypes *in vivo*. Transcriptional profiling and mechanistic studies revealed that HDAC3 represses p65 NF-κB-mediated induction of the Senescence Associated Secretory Program (SASP) and HDAC3 binds directly at the promoters of SASP CXC chemokines. Additionally, HDAC3 was found to cooperate with the lung cancer lineage transcription factor NKX2-1 to mediate expression of a common set of target genes. Leveraging observations about one HDAC3/NKX2-1 common target, FGFR1, we identified that an HDAC3-dependent transcriptional cassette becomes hyperactivated as Kras mutant cancer cells develop resistance to the MEK inhibitor Trametinib, and this can be rescued by treatment with the Class I HDAC inhibitor Entinostat. These unexpected findings reveal new roles for HDAC3 in proliferation control in tumors *in vivo* and identify specific therapeutic contexts for the utilization of HDAC3 inhibitors, whose ability to mechanistically induce SASP may be harnessed therapeutically.

---

Targeted therapies have begun to prove themselves as successful treatments against cancer types harboring specific, defined vulnerabilities. However, only a small subset of tumor types have targeted therapies currently available, as such agents only exist for a limited number of oncogenic drivers. Moreover, tumors characterized by loss of tumor suppressor genes provide no clear targets against which to develop inhibitors. Transcriptional dependencies of tumors have emerged as definable and therapeutically-tractable liabilities that can be oncogene-agnostic (1). Much recent effort has focused on targeting epigenetic regulators (e.g. Brd4) as a means to globally affect transcription in such tumors (2–6). One case in point are Histone Deacetylase (HDAC) inhibitors, which were originally developed to antagonize the reduced global histone acetylation observed in many tumor types (7, 8). Clinically well tolerated, several HDAC inhibitors are now FDA approved to treat blood-borne tumor types (9), although efficacy of HDAC inhibitors in solid tumors has been disappointingly limited. Recent efforts to identify effective approaches to HDAC inhibitor combination therapy have gained traction in specific tumor types (10–14). However, current FDA-approved inhibitors target multiple Class I HDACs, and better therapeutic potential may be realized with more selective inhibitors aimed at one or two HDACs. Despite the fact HDAC inhibitors are already in the clinic, little analysis of Class I HDACs has been performed in genetically engineered solid tumor models in mice.

HDAC3, a Class I HDAC, is unique amongst all HDACs in requiring the Nuclear Receptor Co-Repressor (NCoR) complex for its enzymatic activity (15). HDAC3 has been shown to deacetylate histone and non-histone proteins, and can function in part through deacetylase-independent mechanisms (15). HDAC3 deletion *in vivo* has identified strikingly tissue specific biological functions and associated transcriptional programs (15), revealing that HDAC3 function is not uniformly through global control of histone acetylation, but is nuanced and directed in a tissue-specific fashion. For example, HDAC3 deletion in brown adipose tissue causes mice to become hypothermic and succumb to acute cold exposure (16), but HDAC3 deletion in the liver induces hypertrophy and metabolic alterations (17–19). Despite clinical advancement of inhibitors of Class I HDACs as therapeutics, any potential role of Class I histone deacetylase HDAC3 in cell proliferation and tumorigenesis remains largely unknown, as its *in vivo* function has predominantly been in examined in metabolic tissues.

## HDAC3 is essential for lung tumorigenesis *in vivo*

To assess the role of HDAC3 in solid tumors *in vivo*, we utilized two mouse models engineered to recapitulate the most common subtypes of Kras-mutant Non-Small Cell Lung Cancer (NSCLC); mutant Kras combined with LKB1 loss, Kras^LSL-G12D/+^ STK11^−/−^ (KL), and mutant Kras combined with p53 loss, Kras^LSL-G12D/+^ p53^−/−^ (KP). We first examined mice harboring Kras^LSL-G12D/+^, STK11^L/L^, ROSA26^LSL-luciferase^, with or without conditional HDAC3^L/L^ (KL-HDAC3). In these mice, intratracheal administration of lentivirus expressing Cre recombinase simultaneously activates Kras^G12D^ and deletes LKB1 (*STK11*) to initiate tumorigenesis in the lung epithelium, and for those bearing HDAC3^L/L^, coincidentally deletes HDAC3. Simultaneous induction of firefly luciferase in infected cells allows for noninvasive longitudinal bioluminescence imaging (BLI) of NSCLC tumor development in the whole animal as we have reported previously (20–23). Tumor growth was markedly reduced in KL-HDAC3 mice compared to KL littermate controls at both early and middle timepoints, exhibiting significantly less tumor area, tumor number and smaller tumor size (Fig. 1A-C, S1A-C). Thus, we conclude that HDAC3 supports tumor initiation and tumor growth in the KL model of NSCLC. Employing a similar experimental design, we generated mice harboring Kras^LSL-G12D/+^, p53^L/L^, ROSA26^LSL-luciferase^, HDAC3^L/L^ (KP-HDAC3) to test the role of HDAC3 in the KP model of NSCLC. Tumor growth was dramatically reduced in KP-HDAC3 mice compared to KP littermate controls, with significantly less tumor area and smaller tumor size, and a trend toward smaller tumor number (Fig. 1D-F, S1D). We conclude that HDAC3 is of critical importance for growth of NSCLC tumors driven by both KL and KP genotypes.

**Figure 1.**
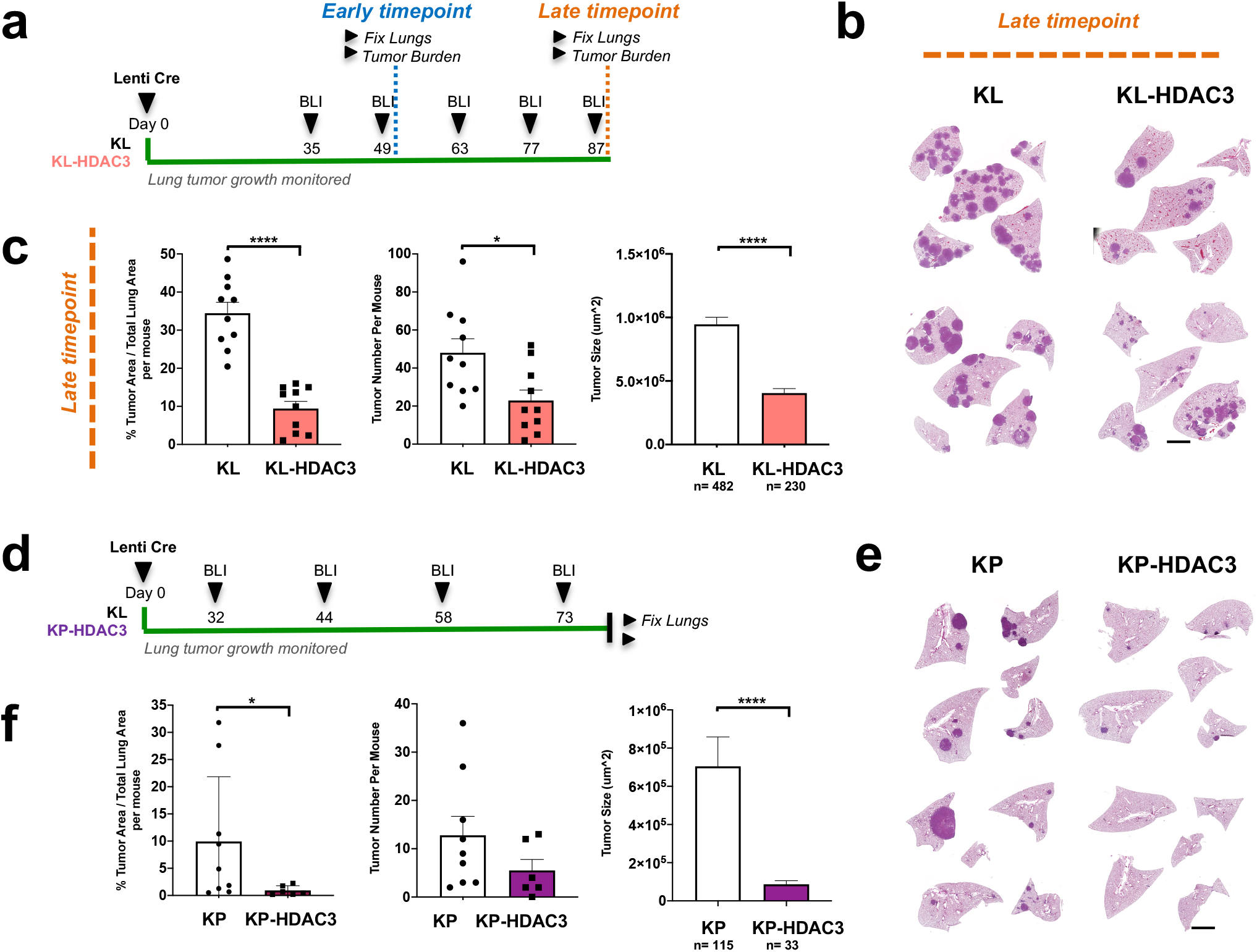
HDAC3 is essential for lung tumorigenesis *in vivo* in KL and KP GEMM models of NSCLC. **(A)** Schematic of experimental design in Kras^G12D/+^, LKB1^L/L^ (KL) and KL-HDAC3^L/L^ (KL-HDAC3) mouse models administered Lenti-Cre. **(B)** Representative H&E-stained sections from the late timepoint. Scale bar 1000um. **(C)** Quantitation from H&E-stained sections from the late timepoint cohort: Tumor area as a percentage of total lung area per mouse (n=10), tumor number per mouse (n=10), and average tumor size (n=482 or 230 as indicated). **(D)** Schematic of experimental design in Kras^G12D/+^, p53^L/L^ (KP) and KP-HDAC3^L/L^ (KP-HDAC3) mouse models administered Lenti-Cre. **(E)** Representative H&E-stained sections. Scale bar 1000um. **(F)** Quantitation from H&E-stained sections: Tumor area as a percentage of total lung area per mouse (n=9 or 6 as indicated), tumor number per mouse (n=9 or 6 as indicated), and average tumor size (n=115 or 33 as indicated). Values are expressed as mean ± s.e.m. * p-value < 0.05, **** p-value < 0.0001 determined by two-tailed Mann-Whitney test.

## HDAC3 represses the p65 NF-κB SASP transcriptional program

To define the mechanisms underlying the anti-tumor growth phenotypes upon HDAC3 deletion, we deleted HDAC3 in KL and KP tumor cell lines derived from the genetically engineered mouse models (GEMM) using CRISPR-Cas9. Cell lines from KL primary tumors are not readily available due to the fact that, unlike KP tumor cells which lack p53, explanted KL primary tumor cells do not grow in culture, presumed to be from p53 activation-dependent growth arrest. To circumvent this issue, we immortalized explanted KL tumor cells before onset of growth arrest. We plucked individual tumors from KL mice and, after dissociation and collagenase treatment, isolated cells were immortalized with SV40 T-antigen and subsequently purified by Epcam+ cell sorting to generate the epithelial lung tumor cell lines KL LJE1 and KL LJE7 (Fig. S2A). These KL cell lines and two established KP cell lines, 634T (24) and KP T3 (20), were infected with lentivirus expressing Cas9 and a gRNA directed against a non-targeting sequence (NT) or HDAC3. Subsequent puromycin selection generated a pooled population of NT or HDAC3 knockout (HDAC3 KO) cells, and immunoblot was used to verify deletion (Fig. 2A, S2B-D). HDAC3 KO reduced cell growth rates across all four cell lines (Fig. S2E), consistent with the anti-tumor growth phenotype of HDAC3 deletion observed *in vivo*.

**Figure 2.**
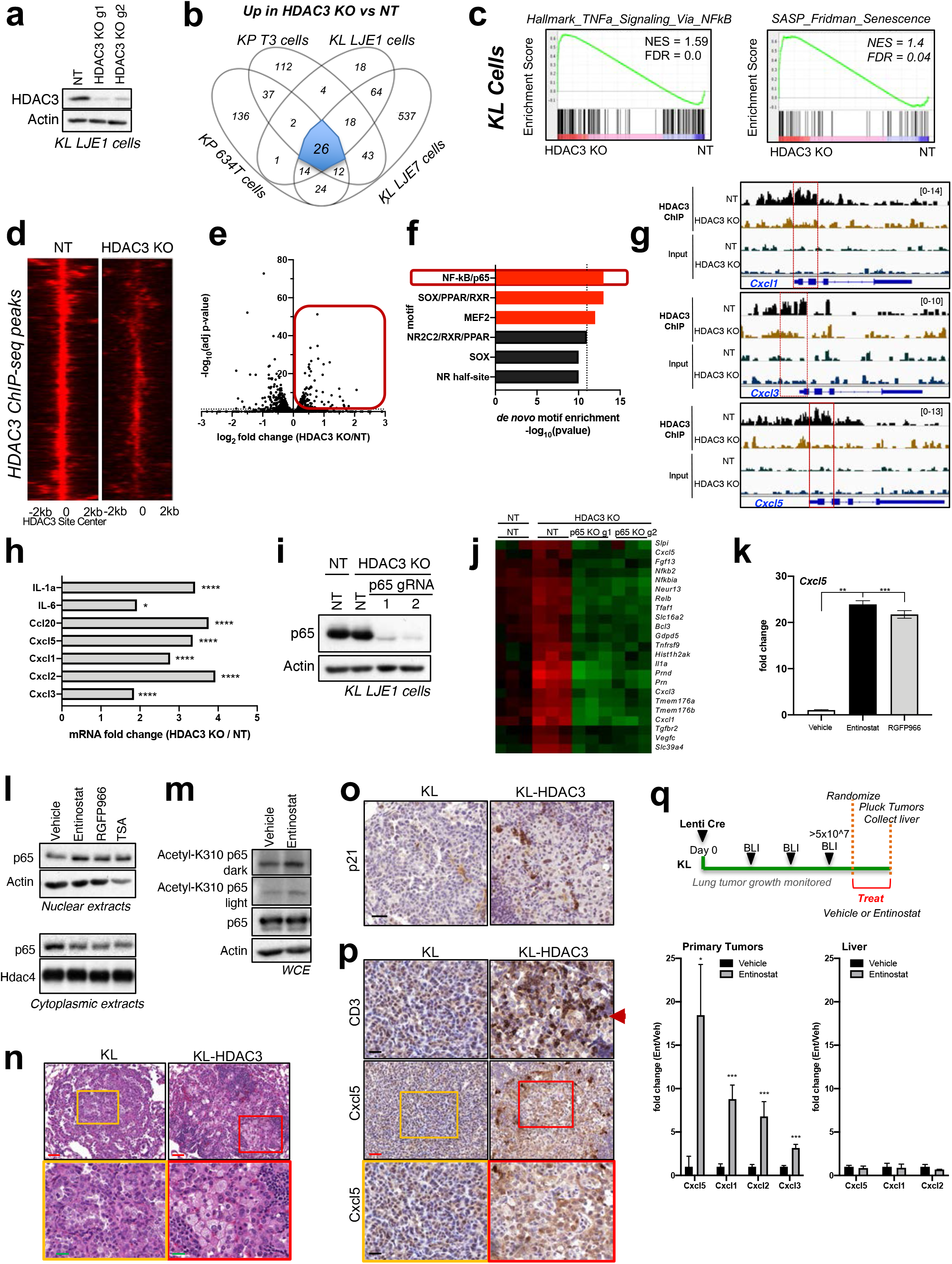
HDAC3 represses the p65 NF-κB SASP transcriptional program in KL and KP NSCLC cells. **(A)** Western blot analysis of HDAC3 deletion (KO) by CRISPR/Cas9 using two different gRNAs (g1, g2) in polyclonal lysates in KL LJE1 cells. **(B)** Overlap of genes upregulated upon HDAC3 KO (using all gRNAs tested) compared to non-targeting (NT) control using RNA-seq data from two KL cells lines (LJE1, LJE7) and two KP cell lines (T3, 634T), with adj. p-value <0.05 and fold change >+/−0.5 cut-offs. **(C)** Gene Set Enrichment Analysis (GSEA) plots of the “Hallmark TNFa Signaling Via NFkB” and “SASP Fridman Senescence” gene sets queried against RNA-seq data comparing HDAC3 KO vs NT conditions across KL LJE1 and KL LJE7 cells combined. **(D)** 3,728 HDAC3 ChIP-seq peaks identified in KL LJE1 NT cells normalized to NT Input and using HDAC3 ChIP-seq in HDAC3 KO cells as background. **(E)** Plot of RNA-seq data from KL LJE1 cells (HDAC3 KO vs NT) for the 468 genes associated with at least one HDAC3 ChIP-seq peak identified in Figure 2D. **(F)** Homer *de novo* motif enrichment analysis of the genomic regions bound by HDAC3 in ChIP-seq for the genes upregulated upon HDAC3 KO (red box) in Figure 2E. Motifs enriched uniquely among upregulated genes are shown. Red bars indicate statistically significant motifs. **(G)** HDAC3 ChIP-seq and Input tracks in NT and HDAC3 KO KL LJE1 cells at the *Cxcl1*, *Cxcl3*, and *Cxcl5* genomic loci. **(H)** Plot of gene expression change (RNA-seq data) upon HDAC3 KO vs NT in KL LJE7 cells for p65-dependent SASP genes. **(I)** Western blot analysis of p65 deletion in HDAC3 KO cells by CRISPR/Cas9 using two different gRNAs in polyclonal lysates in KL LJE1 cells. **(J)** Heatmap showing fragments-per-kilobase-of-transcript-per-million (FPKM) mapped read counts of one gene cluster (Cluster 1) upregulated upon HDAC3 KO in a p65-dependent manner from RNA-seq data generated from cells in Figure 2I. **(K)** qRT-PCR for *Cxcl5* expression in KL LJE1 cells treated with Vehicle, 2uM Entinostat, or 10uM RGFP966 for 72hr (n=3). **(L)** Western blot detecting p65 protein in nuclear or cytoplasmic fractions from KL LJE1 cells after 6hr Vehicle, 2uM Entinostat, 10uM RGFP966, or 0.5uM TSA treatment. **(M)** Western blot analysis of protein lysates from KL LJE1 cells transiently transfected with GFP-p65 and treated 6hr with Vehicle or 2uM Entinostat. **(N)** Representative histopathology images of KL and KL-HDAC3 tumors. Red scale bar 100um, Green scale bar 20um. **(O)** Representative images of p21 IHC in KL and KL-HDAC3 tumors. Scale bar 30um. **(P)** Representative images of CD3 and Cxcl5 IHC in KL and KL-HDAC3 tumors. Black scale bar 20um, red scale bar 40um. **(Q)** *Top.* Schematic of experimental design. *Bottom.* qRT-PCR on primary tumors from KL mice treated 5d with Vehicle or 10mg/kg/day Entinostat by oral gavage. n=6 individual tumors from 3 different mice. Values are expressed as mean ± s.e.m. * p-value < 0.05, ** p-value < 0.01, *** p-value < 0.001 as determined by two-tailed student’s t-test.

To identify the transcriptional programs regulated by HDAC3 in Kras-driven NSCLC cells, we profiled these cell lines by RNA sequencing (RNA-seq). Since HDAC3 KO reduces lung tumor cell growth in KL and KP cells *in vitro* and *in vivo*, we hypothesized that a common mechanism may be engaged upon HDAC3 KO across genotypes, and therefore explored genes commonly deregulated upon HDAC3 KO. Overlap of the RNA-seq datasets across the two KL cell lines and two KP cell lines +/− HDAC3 KO using two different gRNAs identified a stringent set of 38 genes consistently deregulated upon HDAC3 KO in all cell lines profiled (adj. p-value < 0.05, fold > +/−0.5). 26 of these genes were upregulated upon HDAC3 KO (Fig. 2B), and 12 were downregulated upon HDAC3 KO (Fig. S2F). The “Senescence and Autophagy in Cancer” pathway and the “NFKB1” (NF-κB) transcription factor (TF) were the most enriched terms associated with the genes upregulated upon HDAC3 KO (Fig. S2G). The p65/RelA subunit of nuclear factor-κB (NF-κB) is well-established as the TF responsible for inducing the expression of the required Senescence Associated Secretory Phenotype (SASP) gene expression program, engaged as a cell undergoes the largely irreversible program of cell cycle arrest known as senescence (25–28). Gene Set Enrichment Analysis (GSEA) confirmed the upregulation of the SASP Senescence and NF-κB target gene sets upon HDAC3 KO in KL and KP cells (Fig. 2C, S2H).

We next performed HDAC3 ChIP sequencing (ChIP-seq) in NT and HDAC3 KO cells to identify direct HDAC3 targets in KL LJE1 cells (Fig. 2D, S2I). Overlap with the RNA-seq data from KL LJE1 cells revealed that 628 (17%) of the 3,728 HDAC3 peaks were associated with genes differentially expressed upon HDAC3 KO. 364 peaks were associated with 268 genes downregulated upon HDAC3 KO, and 264 peaks were associated with 200 genes upregulated upon HDAC3 KO (Fig. 2e). Motif enrichment analysis of HDAC3-bound genomic regions revealed that the NF-κB/p65 motif was the top *de novo* motif enriched uniquely among genes upregulated upon HDAC3 KO (Fig. 2F), suggesting that in NSCLC lung cancer cells HDAC3 directly binds near p65 and represses p65 target gene expression. We more closely examined HDAC3 binding near several SASP gene loci and found that HDAC3 binds near the Transcription Start Site (TSS) of CXC chemokines in WT but not HDAC3 KO KL cells (Fig. 2G). Importantly, the p65 NF-κB binding elements of each of these genes has been fully annotated and all lie within 100bp of the TSS (29–32). Notably, we did not observe differences in histone acetylation at the HDAC3-bound genomic regions associated with upregulated gene expression (Fig. S2J), suggesting that HDAC3 regulation of these genes is independent of histone acetylation control. Thus, we hypothesized that HDAC3 deletion induces expression of the SASP gene cassette via de-repression of p65.

The SASP genes previously defined to be induced upon engagement of Oncogene-Induced Senescence in a p65-dependent manner (25) exhibit elevated expression levels upon HDAC3 KO in KL cells (Fig. 2H). To test if p65 was required for the induction of these genes following HDAC3 deletion, we knocked out p65 in HDAC3 KO cells using CRISPR/Cas9 (Fig. 2I). qRT-PCR confirmed that the SASP genes *IL-1a*, *Cxcl5*, and *Ccl20* are induced upon HDAC3 KO in a p65-dependent manner (Fig. S2K). RNA-seq on these cells revealed two clusters of genes induced upon HDAC3 KO in a p65-dependent manner (Fig. 2J, S2L), confirming that in NSCLC cells HDAC3 represses the p65 gene expression program which includes SASP genes.

Treatment of KL LJE1 cells with the HDAC inhibitors Entinostat (the clinically viable drug which targets HDAC1 and HDAC3) and RGFP966 (HDAC3 specific) also induced *Cxcl5* gene expression (Fig. 2K), indicating that SASP target genes become induced upon both genetic and pharmacological inhibition of HDAC3. Mechanistically, HDAC3 has been shown to repress NF-κB activity by directly deacetylating p65, which is thought to control its subcellular localization (33–35), although the engagement of this molecular mechanism *in vivo* has not been widely reported. Treatment of KL LJE1 cells for 6 hours with HDAC inhibitors Entinostat, RGFP966, and TSA (pan-HDAC inhibitor) induce accumulation of endogenous p65 protein in the nuclear compartment and corresponding depletion of p65 in the cytoplasm (Fig. 2L). To facilitate detection of one of the previously reported acetylation sites on p65, lysine 310 (33), we next transfected KL LJE1 cells with GFP-tagged p65 (Fig. 2M) or Flag-tagged p65 (Fig. S2M). 6 hours of Entinostat treatment induced acetylation of K310 on p65 as detected by western blot of protein lysates. Repression of NF-κB function by HDAC3 has been observed in macrophages (36), but has not been reported in the many other HDAC3-tissue specific knockouts. We have identified that in both KL and KP NSCLC cells, HDAC3 actively represses the p65 NF-κB transcriptional program, resulting in repression of the tumor cell SASP program. Importantly, this suggests that HDAC3-targeting inhibitors can be employed to modulate p65 activity and the senescence program.

To explore if HDAC3 is repressing p65 and the SASP senescence program *in vivo*, we analyzed histological sections from KL-HDAC3 and KP-HDAC3 mice (Fig. 1). Analysis of H&E sections revealed that KL-HDAC3-deleted tumors exhibit unusual epithelial cell morphology characterized by enlarged cytoplasm (Fig. 2N). Immunohistochemistry (IHC) for senescence marker p21 revealed that HDAC3 deleted tumors exhibit elevated levels of this marker (Fig. 2O), consistent with increased rates of senescence upon HDAC3 deletion *in vivo*. Furthermore, KL-HDAC3 and KP-HDAC3 tumors displayed aberrant recruitment of immune cells to the periphery of and infiltrating the tumor compared to wildtype controls (Fig. 2N, S2N). Based on our molecular model, we hypothesized that the HDAC3-dependent immune cell recruitment could be due to de-repression of the SASP program, and sought to determine the identity of the recruited immune cells. IHC identified that CD3+ T-cells are present within KL and KP tumors deleted for HDAC3 to a greater extent than in wildtype tumors of either genotype (Fig. 2P, S2O). F4/80+ macrophages, NKp46+ NK cells, and Ly6g+ neutrophils did not exhibit this HDAC3-dependent pattern (Fig. S2P). We reasoned that if this T-cell recruitment was due to p65/NF-κB-driven induction of the SASP program, p65 target gene expression should be elevated in HDAC3 deleted in tumors. Consistently, IHC identified elevated levels of Cxcl5 in HDAC3 KO tumors (Fig. 2P, S2O), a key SASP target gene that is among the core genes consistently transcriptionally deregulated in an HDAC3- and p65-dependent manner (Fig. 2B, H, J). Thus, HDAC3 is an *in vivo* regulator of the SASP cellular senescence program in lung tumors.

To determine if HDAC inhibitors induce SASP target gene expression in primary tumors *in vivo*, we next treated lung tumor-bearing KL mice with Vehicle or 10mg/kg/day Entinostat for 5 days by oral gavage, and subsequently collected lung tumors and livers (Fig. 2Q). RNA was isolated from treated tissues, and qRT-PCR determined that Entinostat induces expression of SASP target genes *Cxcl5, Cxcl1, Cxcl2,* and *Cxcl3* in primary KL tumors *in vivo*. Importantly, expression of these genes was not induced by Entinostat in the livers of these mice, suggesting a tumor-specific effect.

## HDAC3 cooperates with NKX2-1 to regulate the expression of a common set of target genes

To further define the critical HDAC3 targets in tumorigenesis, HDAC3 ChIP-seq was performed on KL and KP primary tumors to identify genome wide, endogenously bound HDAC3 target loci *in vivo* (Fig. S3A-B, 3A). 1522 peaks were bound by HDAC3 in both KL and KP tumors (Fig. 3A), corresponding to 753 non-redundant genes with at least one HDAC3 binding site within +/− 25 kilobases of the TSS. We queried the expression pattern of these 753 genes across individual primary tumors isolated from four different Kras-driven lung tumor GEMMs; Kras, KP, KL, and KPL (Fig. 3B). A large fraction of the 753 HDAC3-bound genes exhibited an LKB1-dependent transcriptional pattern, suggesting that an LKB1-dependent mechanism may be mediating differential regulation of a subset of these target genes bound by HDAC3 in both KL and KP tumors. Comparing gene expression between KL versus Kras primary tumors, we found that 39% of the 753 HDAC3 target genes were differentially expressed upon LKB1 loss (Fig. S3C). Motif enrichment analysis of the HDAC3-bound sites common to KL and KP tumors (Fig. 3A) identified the top enriched *de novo* motif to be that of the transcription factor NKX2-1 (TTF-1) (Fig. 3C), a clinical biomarker of lung adenocarcinoma which has an established role enforcing a lineage-specific differentiation program in lung and lung ADC (37–40). NKX2-1 is an appreciated but undruggable transcriptional addiction of LUAD. Therefore, identifying druggable regulators of NKX2-1 function is of great interest. While the p65 motif was also identified in this analysis, its enrichment was lower than that of the NKX2-1 motif, suggesting that HDAC3 may interact with p65 at a smaller subset of target genes, whereas HDAC3 interplay with NKX2-1 may occur at a broader set of targets. This analysis suggested an unexpected functional overlap between HDAC3 and NKX2-1, and co-immunoprecipitation experiments confirmed that endogenous NKX2-1 interacts with HDAC3 in KL LJE1 cells (Fig. 3D).

**Figure 3.**
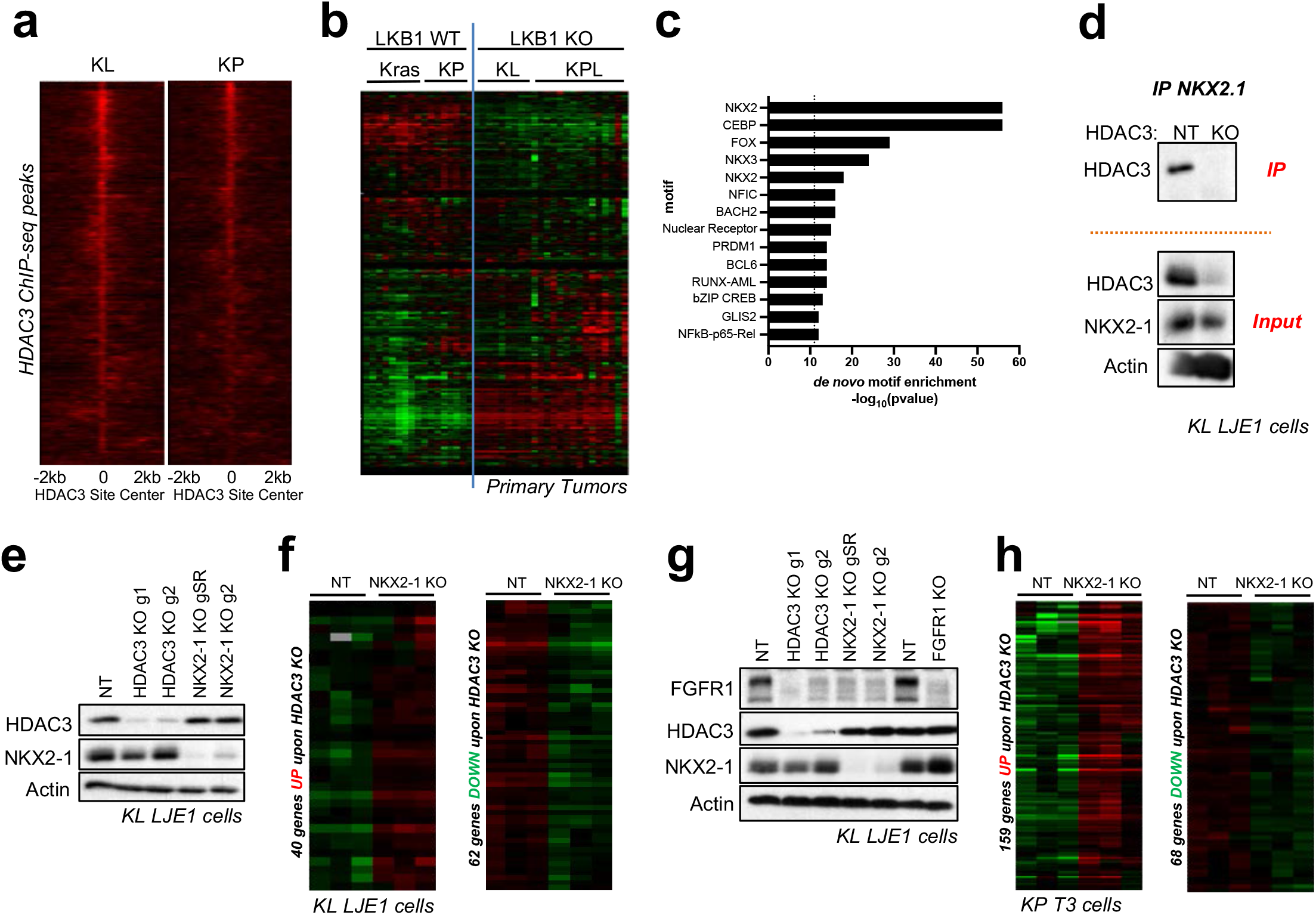
HDAC3 cooperates with NKX2-1. **(A)** 1522 HDAC3 ChIP-seq peaks common to KL and KP primary tumors. **(B)** Heatmap of RNA-seq data showing FPKM read counts from primary tumors from LKB1 WT (Kras, KP) and LKB1 KO (KL, KPL) models for the 753 non-redundant genes associated with at least one HDAC3 ChIP-seq peak within 25kb of the TSS. **(C)** Homer *de novo* motif enrichment analysis of the HDAC3-bound peaks in Figure 3A. All significantly enriched motifs are listed. **(D)** Western blot analysis of NKX2-1 co-immunoprecipitation with HDAC3 in NT vs HDAC3 KO KL LJE1 cells. **(E)** Western blot analysis of HDAC3 and NKX2-1 deletion (KO) by CRISPR/Cas9 using two different gRNAs (gSR, g2 for NKX2-1) in polyclonal lysates in KL LJE1 cells. **(F)** Heatmap of RNA-seq data showing FPKM read counts for genes commonly upregulated (left) or downregulated (right) upon both HDAC3 KO and NKX2-1 KO in KL cells, as defined from red box regions on heatmap in Figure S4G. **(G)** Western blot analysis of FGFR1, HDAC3 and NKX2-1 in NT, HDAC3 KO, NKX2-1 KO, and FGFR1 KO cell lysates from KL LJE1 cells. **(H)** Heatmap of RNA-seq data showing FPKM read counts for genes commonly upregulated (left) or downregulated (right) upon both HDAC3 KO and NKX2-1 KO in KP T3 cells, as defined from red box regions on heatmap in Figure S4J.

To explore if NKX2-1 and HDAC3 co-regulate a set of transcriptional targets, we used CRISPR/Cas9 to delete NKX2-1 in KL LJE1 cells (Fig. 3E), and profiled these cells by RNA-seq. Plotting gene expression in KL LJE1 NT or HDAC3 KO cells for the genes most de-regulated upon NKX2-1 KO revealed that 15/18 (83%) are also modulated upon HDAC3 KO (Fig. S3D), revealing that the most NKX2-1-dependent genes are nearly all under the control of HDAC3. To query the extent to which HDAC3 is involved in the regulation of NKX2-1 target genes across a broader set of genes, we extended this analysis to the 68 genes deregulated upon NKX2-1 KO by the stronger gRNA (SR), and found that 72% of NKX2-1 target genes were also modulated upon HDAC3 KO in these cells (data not shown) and 31% were associated with HDAC3 ChIP-seq peaks. This suggests that nearly three quarters of the entire NKX2-1 transcriptional program is co-regulated by HDAC3. We found that of the 28 genes deregulated upon NKX2-1 KO and HDAC3 KO, 21/22 (96%) were downregulated and 6/6 (100%) were upregulated upon loss of both NKX2-1 and HDAC3 (Fig. S3E). Thus, HDAC3 KO and NKX2-1 KO affect gene expression changes with the same directionality. Consistently, using previously published gene expression data from Kras^mut^ tumors tamoxifen-induced to delete NKX2-1 (37), we found that genes downregulated upon NKX2-1 in Kras^mut^ tumors were negatively enriched upon HDAC3 KO, and genes upregulated upon NKX2-1 loss in Kras^mut^ tumors were positively enriched for upon HDAC3 KO in KL LJE1 cells (Fig. S3F).

To assess what fraction of the total HDAC3 transcriptional response is regulated by NKX2-1, we plotted gene expression from NT and NKX2-1 KO cells for all genes differentially expressed upon HDAC3 KO in both KL LJE1 and KL LJE7 cells (Fig. 3F, S3G,). Of the 171 genes upregulated upon HDAC3 KO, 21% were also upregulated but only 3% downregulated upon NKX2-1 KO. Of the 165 genes downregulated upon HDAC3 KO, 38% were also downregulated and none upregulated upon NKX2-1 KO. Notably, amongst those genes regulated in a common direction upon HDAC3 or NKX2-1 loss (up vs down), HDAC3 ChIP-seq peaks were found associated with genes both up- (25%) and down- (37%) regulated indicating that unlike p65 NF-κB SASP genes, HDAC3 is not simply acting to repress the NKX2-1 controlled genes.

One co-regulated target gene of NKX2-1 and HDAC3 (Fig. 3F) in KL LJE1 cells is *Fgfr1*. Fgfr1, Fibroblast Growth Factor Receptor 1, is one of four receptor tyrosine kinases that make up the FGFR protein family. FGFRs are receptors for Fibroblast Growth Factors (FGFs), have been widely implicated in promoting tumor growth, and multiple small molecular inhibitors of FGFRs are in various stages of development as cancer therapies (41, 42). We found that *Fgfr1* mRNA was downregulated upon both NKX2-1 KO and HDAC3 KO, and *Fgfr1* was associated with an HDAC3 ChIP-seq peak. Western blotting identified robust reduction in FGFR1 protein level upon HDAC3 KO or NKX2-1 KO in KL LJE1 cells (Fig. 3G), confirming that HDAC3 and NKX2-1 coordinately regulate FGFR1 in KL cells.

To explore FGFR1 regulation in KP cells, we used CRISPR/Cas9 to delete NKX2-1 in KP T3 cells. Surprisingly, we found that neither HDAC3 KO nor NKX2-1 KO altered FGFR1 protein levels in KP T3 cells (Fig. S3H), suggesting differential engagement of the HDAC3/NKX2-1 complex in the regulation of this target in KL versus KP cells. To query if this was unique to FGFR1 - or if FGFR1 was an indication of broader functional differences between HDAC3/NKX2-1 in KL versus KP cells - we profiled KP T3 cells deleted for HDAC3 or NKX2-1 (Fig. S3H) by RNA-seq. Performing analysis on data from KP T3 cells comparable to that done on data from KL LJE1 cells (Fig. 3F, S3D, G) revealed that HDAC3 and NKX2-1 coordinately *repress* the expression of a set of common target genes in KP T3 cells (Fig. S3I-J, 3H). Of the 202 NKX2-1-dependent genes, 74% were deregulated by HDAC3 loss in KP cells, and 91% of these genes were upregulated, demonstrating that in KP cells NKX2-1 and HDAC3 loss results in enhanced gene expression of shared target genes. Of the 388 HDAC3-dependent genes in KP T3 cells, one third were downregulated and two thirds were upregulated upon HDAC3 loss (Fig. S3J). 63% of genes upregulated upon HDAC3 KO were also upregulated upon NKX2-1 KO (Fig. S3J, 3H), whereas NKX2-1 KO did not strongly modulate expression of genes downregulated upon HDAC3 KO. This suggests that HDAC3 and NKX2-1 cooperate to <u>repress</u> a set of common target genes in KP T3 cells. This is in contrast with KL LJE1 cells, where HDAC3 and NKX2-1 <u>promote</u> the expression of a set of common target genes. Consistent with HDAC3 and NKX2-1 cooperating to regulate divergent gene programs in KL versus KP cells, little overlap was observed between the co-regulated genes in KL LJE1 and KP T3 cells (Fig. S3K). Notably, HDAC3-mediated repression of p65 target gene expression was observed consistently across these same KL and KP lung tumor cell lines regardless of genotype (Fig. 2). In contrast to HDAC3 repression of p65, the data regarding NKX2-1 raises the hypothesis that HDAC3 and NKX2-1 may co-regulate different gene programs depending on the tumor genotype.

## HDAC3 and NKX2-1 co-regulate target genes which are aberrantly engaged upon Trametinib resistance

Since FGFR1 was regulated by HDAC3 and NKX2-1 in our Kras-mutant LUAD cells in a genotype-dependent manner, we utilized it as an example with which to explore the divergence of HDAC3/NKX2-1 co-regulatory function between KL and KP cells. We found that, in primary tumors, *Fgfr1* mRNA is elevated in LKB1 KO compared to LKB1 WT tumors (Fig. S4A). Deletion of FGFR1 with CRISPR/Cas9 confirmed that FGFR1 KO reduced KL LJE1 cell growth rates (Fig. S4B), supporting a role for FGFR1 in sustaining tumor cell growth. In The Cancer Genome Atlas (TCGA) Lung Adenocarcinoma dataset, high *FGFR1* mRNA correlated with poor patient outcome and, interestingly, 25% of the *FGFR1*-high tumors harbor *STK11* (LKB1) mutation (Fig. S4C).

Notably, FGFR1 has been shown to mediate resistance to the FDA-approved MEK inhibitor, Trametinib, that acts downstream of Kras to suppress signaling through the mitogen-activated protein kinase (MAPK) cascade (43). Manchado et al. found that co-treatment of Trametinib with the FGFR1 inhibitor Ponatinib induced tumor regression in the KP lung tumor model where Trametinib treatment alone is largely ineffective, due to rapidly acquired resistance (43). Despite the high rate of Kras mutation in LUAD (~30%), therapeutic targeting of the Kras pathway remains a major challenge in part because therapies directed against Kras effectors activate compensatory pathways that limit their efficacy as single agents. Many current efforts are directed toward elucidating which combination therapy approaches would potentiate clinical benefit from existing Kras effector inhibitors.

Since HDAC3 supports FGFR1 protein expression in KL LJE1 cells (Fig. 3G), we hypothesized that HDAC inhibition may be an alternative approach to blocking the induction of FGFR1 by long-term Trametinib treatment in lung cancer cells. We therefore performed short-term (3 day) and long-term (13 day) treatments with Vehicle or Trametinib. In KL LJE1 cells, FGFR1 protein was strongly induced upon 13 day Trametinib in a manner that could be rescued by co-treatment of Trametinib plus Entinostat, a clinically well-tolerated Class I HDAC inhibitor (Fig. 4A). This implies that HDAC inhibitors which target HDAC3 (such as Entinostat) can block the induction of a transcriptional program that is engaged upon Trametinib resistance. Interestingly in KP T3 cells, 13 day Trametinib treatment did not induce FGFR1 levels (Fig. S4D).

**Figure 4.**
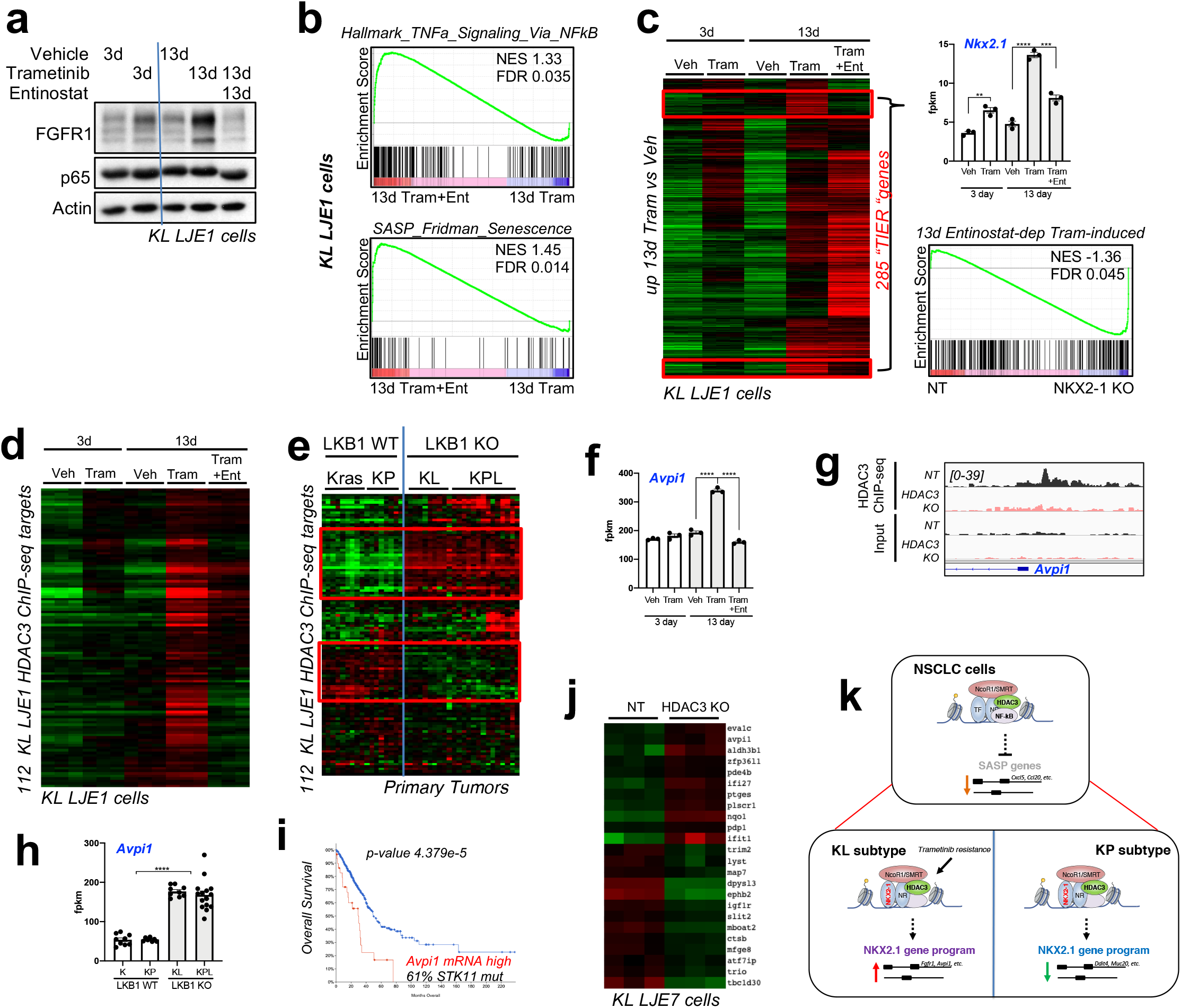
HDAC3 and NKX2-1 regulate the expression of a common set of target genes which is aberrantly engaged upon Trametinib resistance. **(A)** Western blot analysis of FGFR1 and p65 in protein lysates from KL LJE1 cells treated with Veh, 10nM Trametinib, or 1uM Entinostat for 3 or 13 days. **(B)** GSEA plots of the “Hallmark TNFa Signaling Via NFkB” and “SASP Fridman Senescence” gene sets queried against RNA-seq data comparing 13 day Trametinib+Entinostat to 13 day Trametinib conditions in KL LJE1 cells. **(C)** *Left:* Heatmap of RNA-seq data showing FPKM read counts across all treatment conditions for the 2,141 genes significantly upregulated (adj. p-value < 0.05, fold > +/−0.5) upon 13 day Trametinib compared to 13 day Vehicle in KL LJE1 cells. Veh, Vehicle; Tram, Trametinib; Ent, Entinostat. *Right, top: Nkx2-1* mRNA levels across all treatment conditions (n=3). *Right, bottom:* GSEA analysis of the 285 Trametinib-induced, Entinostat-rescued (“TIER”) genes queried across RNA-seq data from NKX2-1 KO vs NT KL LJE1 cells. **(D)** Heatmap of RNA-seq data showing FPKM read counts across all treatment conditions for the 112 TIER genes which are HDAC3 ChIP-seq target genes. Veh, Vehicle; Tram, Trametinib; Ent, Entinostat. **(E)** Heatmap of RNA-seq data showing FPKM read counts from Kras (n=9), KP (n=8), KL (n=9), and KPL (n=15) primary tumors for the 112 TIER genes which are HDAC3 ChIP-seq target genes (Figure 4D). **(F)** *Avpi1* mRNA levels across all treatment conditions (n=3). Veh, Vehicle; Tram, Trametinib; Ent, Entinostat. **(G)** HDAC3 ChIP-seq data in NT and HDAC3 KO KL LJE1 cells at the *Avpi1* genomic locus. **(H)** *Avpi1* mRNA expression across Kras (K) (n=9), KP (n=8), KL (n=9), and KPL (n=15) primary tumor RNA-seq data (n=8-15). **(I)** Overall survival data for all patients comparing tumors with or without high *AVPI1* mRNA from the Firehose Legacy LUAD TCGA dataset. **(J)** Heatmap of RNA-seq data showing FPKM read counts for the 24 genes within the published 122 gene Nanostring LKB1-mutant NSCLC signature (28) which are deregulated at the mRNA level in HDAC3 KO cells. **(K)** Model of HDAC3 function in Kras-mutant NSCLC cells. Values are expressed as mean ± s.e.m. ** p-value < 0.01, *** p-value < 0.001, **** p-value <0.0001 as determined by two-tailed student’s t-test.

To test if FGFR1 is part of a larger HDAC3-dependent transcriptional cassette that becomes hijacked upon Trametinib resistance in lung cancer cells, we profiled KL LJE1 and KP T3 cells treated as in Fig. 4A and S4D by RNA-seq. First, we found that Entinostat treatment induced expression of the same “TNFa Signaling Via NFkB” and “SASP Fridman Senescence” gene sets (Fig. 4B, S4E) which enriched upon genetic deletion of HDAC3 (Fig. 2C, S2H) in both KL LJE1 and KP T3 cells. Importantly, this confirms that HDAC3 functions as an upstream repressor of the p65 NF-κB and SASP programs in both KL and KP cells in a manner which can be therapeutically targeted using Entinostat, even in the context of Trametinib resistance. However, 13 day Trametinib treatment compared to Vehicle did not enrich for these gene sets, indicating that Trametinib alone does not induce NF-κB repression. Thus, while Entinostat robustly activates NF-κB/SASP in the presence of Trametinib resistance, this mechanism does not appear to be unique to Trametinib resistant cells.

Based on the behavior of FGFR1, we were also interested in looking for NKX2-1 and HDAC3 common targets that become induced upon 13 day Trametinib treatment in a manner reversed by Entinostat. To identify the genes behaving in this pattern, we first defined the genes upregulated upon 13 day Trametinib compared to 13 day Vehicle in KL LJE1 cells, and then plotted the gene expression across all conditions for these 2,141 genes (Fig. 4C). The expression of 285/2141 (13%) genes was uniquely induced upon 13 day Trametinib (not upon 3 day Trametinib) in a manner reversed upon co-treatment of Entinostat. One of these 285 Trametinib-induced, Entinostat-reversed (“TIER”) genes was *Nkx2-1* itself (Fig. 4C, top right), unbiasedly confirming that NKX2-1 induction is a component of long-term Trametinib resistance in KL cells that can be reversed by HDAC inhibition.

To explore if a broader set of NKX2-1-dependent genes are behaving similarly, we queried the 285 TIER genes against RNA-seq data from NKX2-1 KO cells using GSEA analysis (Fig. 4C, bottom right). Indeed, the TIER gene set was negatively enriched upon NKX2-1 KO in basal conditions. We hypothesized that Entinostat reverses the induction of NKX2-1 target gene expression through inhibition of HDAC3, and thus predicted that a set of HDAC3 direct target genes would also behave in a Trametinib- and Entinostat-dependent manner. Of the 2,933 unique genes associated with at least one HDAC3 ChIP-seq binding site, 112 genes were induced upon 13 day Trametinib treatment in a manner rescued by Entinostat (Fig. 4D). We plotted gene expression from Kras, KP, KL, and KPL primary GEMM lung tumor RNA-seq for these 112 direct HDAC3 bound TIER genes and found that 43 of these genes (38%) are expressed in an LKB1-dependent manner (Fig. 4E, S4F). Together, we have identified a set of direct HDAC3 target genes which display LKB1-dependent gene expression in primary tumors, and whose expression is induced in tumor cells as a component of targeted therapy resistance in a manner that can be rescued by HDAC inhibition.

To explore the expression of these genes in human tumors, we queried the TCGA human Lung Adenocarcinoma dataset (Firehose Legacy). We found that one of the most LKB1-dependent of these 43 HDAC3 target genes, *AVPI1* (Fig. 4F-H, S4F), was highly expressed (EXP>1.8) in 30/586 (5%) TCGA human tumor samples. These 30 patients had significantly poorer overall survival (30.29 versus 49.8 median months overall) (Fig. 4I) and time disease free (data not shown). 11/18 (61.1%) *AVPI1*-high tumors with mutational data harbored *STK11* mutation, the most frequently mutated gene in this group, while only 29/212 (13.7%) of non-*AVPI1*-high tumors had *STK11* mutation. 8/18 (44%) *AVPI1*-high tumors also exhibited Kras amplification or mutation at G12, and these patients had even shorter overall survival when compared to the 92 patients with Kras amplified or G12 mutated tumors without high *AVPI1* (8.48 versus 88.07 median months overall) (Fig. S4G). Of *AVPI1*-high, Kras-mutant tumors with mutational data, 5/7 (71.43%) harbored *STK11* mutation and 1/7 had deep deletion of *STK11*. Interestingly, *AVPI1* has been identified from human NSCLC datasets as one of the LKB1 classifier genes whose expression can predict LKB1 mutation in lung adenocarcinoma (44, 45). Together, this suggests that our HDAC3 molecular signature datasets can predict patients destined for poor outcome and those who harbor LKB1 mutant tumors. Further analysis of the 122 genes in a Nanostring LKB1-mutant NSCLC signature (28) revealed that 24 of these genes (20%) are deregulated at the mRNA level in HDAC3 KO cells, half of which exhibit direct HDAC3 binding nearby (Fig. 4J).

Taken altogether, we have found here a critical role for the Class I HDAC3 in lung tumorigenesis and revealed unexpected roles for HDAC3 in restraining cellular senescence and SASP in Kras mutant lung tumors *in vivo*. The direct molecular mechanism we have detailed, with HDAC3 directly binding near the TSSs of SASP gene loci which bear previously mapped p65 NF-κB binding sites in their proximal promoter, suggests a model by which HDAC3 is directly controlling p65 acetylation and target gene expression in these tumors, indicating that induction of SASP genes in tumor cells may be tunable via administration of HDAC3 inhibitors. An additional unexpected finding here is that HDAC3 mediates expression of a substantial fraction of LKB1-mutant tumor specific gene expression. Furthermore, a large percentage of HDAC3-dependent genes are also dependent on the lineage specific transcription factor NKX2-1, which appears to be induced in KL LUAD cells as a resistance mechanism to MEK inhibitors, an effect reversed by HDAC3 inhibition, revealing a specific therapeutic context where HDAC3 inhibition may have great utility (Fig. 4K). These findings motivate further exploration of the role of HDAC3 in epithelial tumors, senescence, and resistance to targeted therapies.

**Correspondence and requests for materials** should be addressed to R.J.S.

---

**Figure S1.**
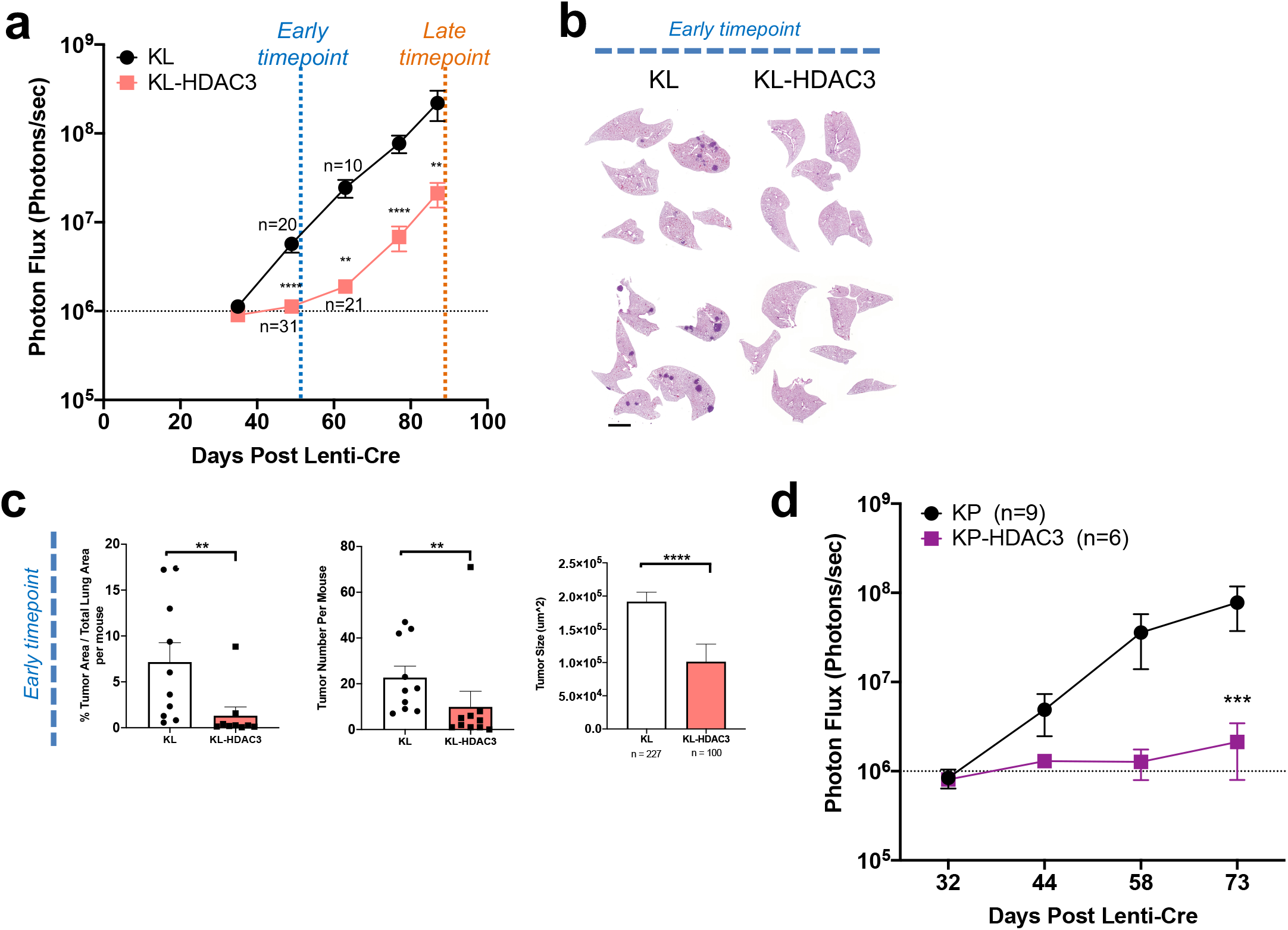
HDAC3 deletion *in vivo* impairs tumor growth in KL and KP GEMM models of NSCLC. **(A)** Average longitudinal BLI data from the KL-HDAC3 experiment. **(B)** Representative H&E-stained sections from the KL-HDAC3 early timepoint. Scale bar 1000um. **(C)** Quantitation from H&E-stained sections from the early timepoint cohort: Tumor area as a percentage of total lung area per mouse (n=10), tumor number per mouse (n=10), and average tumor size (n=227 or 100 as indicated). **(D)** Average longitudinal BLI data from the KP-HDAC3 experiment. Values are expressed as mean ± s.e.m. ** p-value < 0.01, *** p-value < 0.001, **** p-value < 0.0001 determined by two-tailed Mann-Whitney test.

**Figure S2.**
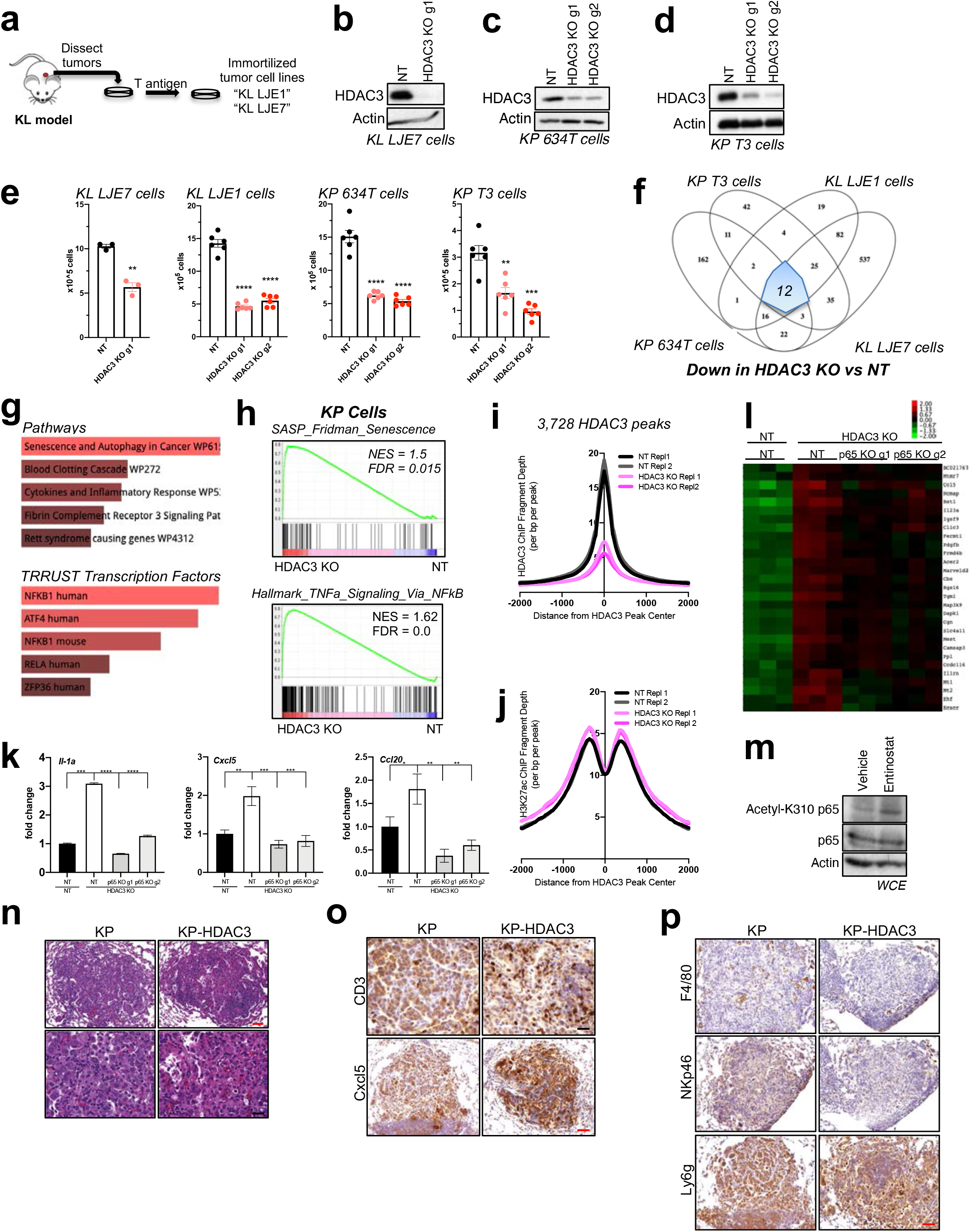
HDAC3 represses the p65 NF-κB SASP transcriptional program in KL and KP NSCLC cells. (A) Schematic of experimental design to establish KL LJE1 and KL LJE7 cell lines from KL primary tumors. (B) Western blot analysis of HDAC3 KO by CRISPR/Cas9 in polyclonal lysates in KL LJE7 cells. (C) Western blot analysis of HDAC3 KO by CRISPR/Cas9 in polyclonal lysates in 634T cells. (D) Western blot analysis of HDAC3 KO by CRISPR/Cas9 in polyclonal lysates in KP T3 cells. (E) Proliferation assessment 5 days after plating HDAC3 KO or NT cells from KL LJE7, KL LJE1, KP 634T, and KP T3 cell lines (n=6). (F) Overlap of genes downregulated upon HDAC3 KO (using all gRNAs tested) compared to NT using RNA-seq data from KL LJE1, KL LJE7, KP T3, and KP 634T cell lines, with adj. p-value <0.05 and fold change >+/−0.5 cut-offs. (G) Enrichr Pathway and Transcription analysis of the 26 commonly upregulated genes identified in Figure 2B. (H) GSEA plots of the “Hallmark TNFa Signaling Via NFkB” and “SASP Fridman Senescence” gene sets queried against RNA-seq data comparing HDAC3 KO vs NT conditions across KP cells. (I) Average HDAC3 ChIP-seq fragment depth +/−2kb of each peak center for the 3,728 HDAC3 ChIP-seq peaks identified in KL LJE1 NT cells in Figure 2D. Data from each ChIP replicate (Repl) from NT and HDAC3 KO cells is plotted. (J) Average H3K27ac ChIP-seq fragment depth +/−2kb of each peak center for the HDAC3 ChIP-seq peaks associated with upregulated gene expression upon HDAC3 KO (red box, Figure 2E) in KL LJE1 NT cells. Data from each ChIP replicate (Repl) from NT and HDAC3 KO cells is plotted. (K) qRT-PCR for *Il-1a*, *Cxcl5*, and *Ccl20* expression in KL LJE1 cells deleted for HDAC3 +/− p65 KO (n=3). (L) Heatmap showing FPKM read counts of the second gene cluster (Cluster 2) upregulated upon HDAC3 KO in a p65-dependent manner from RNA-seq data generated from cells in Figure 2I. (M) Western blot analysis of protein lysates from KL LJE1 cells transiently transfected with Flag-p65 and treated 6hr with Vehicle or 2uM Entinostat. (N) Representative histopathology images of KP and KP-HDAC3 tumors. Red scale bar 50um, black scale bar 20um. (O) Representative images of CD3 and Cxcl5 IHC in KP and KP-HDAC3 tumors. Black scale bar 20um, Red scale bar 50um. (P) Representative images of F4/80, NKp46, and Ly6g IHC in KP and KP-HDAC3 tumors. Scale bar 50um. Values are expressed as mean ± s.e.m. * p-value < 0.05, ** p-value < 0.01, *** p-value < 0.001, **** p-value < 0.0001 determined by two-tailed student’s t-test.

**Figure S3.**
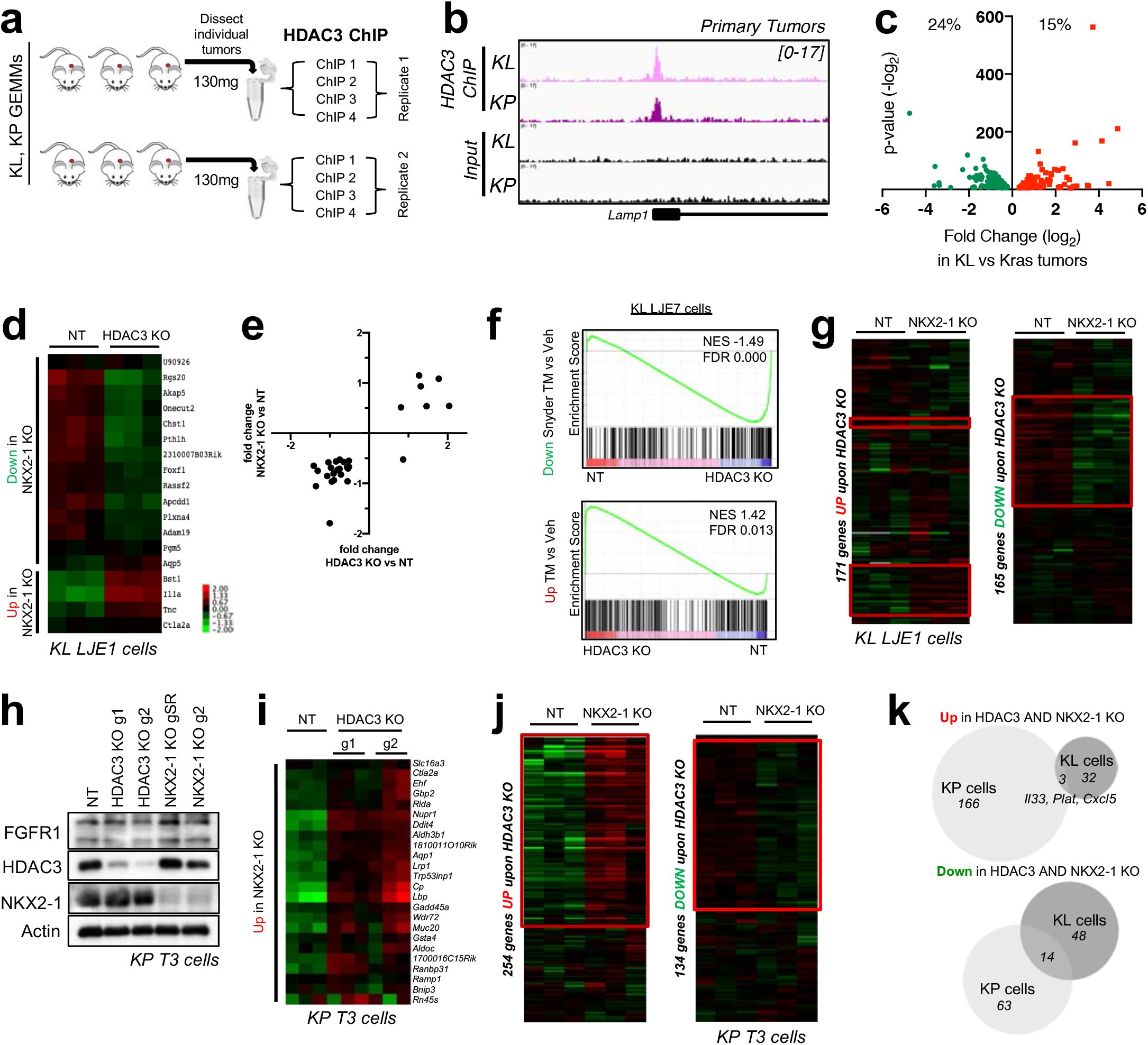
HDAC3 cooperates with NKX2-1. (A) Schematic of HDAC3 ChIP-seq experimental design in primary KL and KP tumors. (B) Example of an HDAC3 ChIP-seq peak at a genomic region bound by HDAC3 in both KL and KP primary tumors. (C) Plot of RNA-seq differential expression between KL versus Kras primary tumors for the HDAC3 target genes in Figure 3A. (D) Heatmap of RNA-seq data showing FPKM read counts from NT or HDAC3 KO cells for the genes significantly deregulated upon NKX2-1 KO (adj. p-value <0.05, fold +/−0.5) in KL LJE1 cells. (E) Plot of fold change upon HDAC3 KO compared to NKX2-1 KO for the genes in Figure S4D. (F) GSEA plots for genes deregulated (downregulated, top plot; or upregulated, bottom plot) upon tamoxifen (TM)-mediated *in vivo* deletion of NKX2-1 in Kras tumors (Snyder et al. Mol Cell, 2013) queried across HDAC3 KO RNA-seq data from KL LJE7 cells. (G) Heatmap of RNA-seq data showing FPKM read counts from NT or NKX2-1 KO KL LJE1 cells for the genes significantly deregulated upon HDAC3 KO (adj. p-value <0.05, fold +/−0.5) in KL LJE1 and KL LJE7 cells. (H) Western blot analysis of FGFR1, HDAC3 and NKX2-1 in NT, HDAC3 KO and NKX2-1 KO cell lysates from KP T3 cells. (I) Heatmap of RNA-seq data showing FPKM read counts from NT or HDAC3 KO cells for the genes significantly deregulated upon NKX2-1 KO (adj. p-value <0.05, fold +/−0.5) in KP T3 cells. 20/24 (83%) of the most stringent NKX2-1-dependent genes were also HDAC3-dependent, and all were upregulated upon deletion of NKX2-1 or HDAC3. (J) Heatmap of RNA-seq data showing FPKM read counts from NT or NKX2-1 KO KL LJE1 cells for the genes significantly deregulated upon HDAC3 KO (adj. p-value <0.05, fold +/−0.5) in KP T3 cells. 159/254 genes were strongly upregulated upon HDAC3 KO or NKX2-1 KO (red box, left), whereas 68/134 genes were mildly downregulated upon HDAC3 KO or NKX2-1 KO (red box, right). (K) Overlap of genes significantly upregulated (top) or downregulated (bottom) upon both HDAC3 KO and NKX2-1 KO in KP versus KL cells.

**Figure S4.**
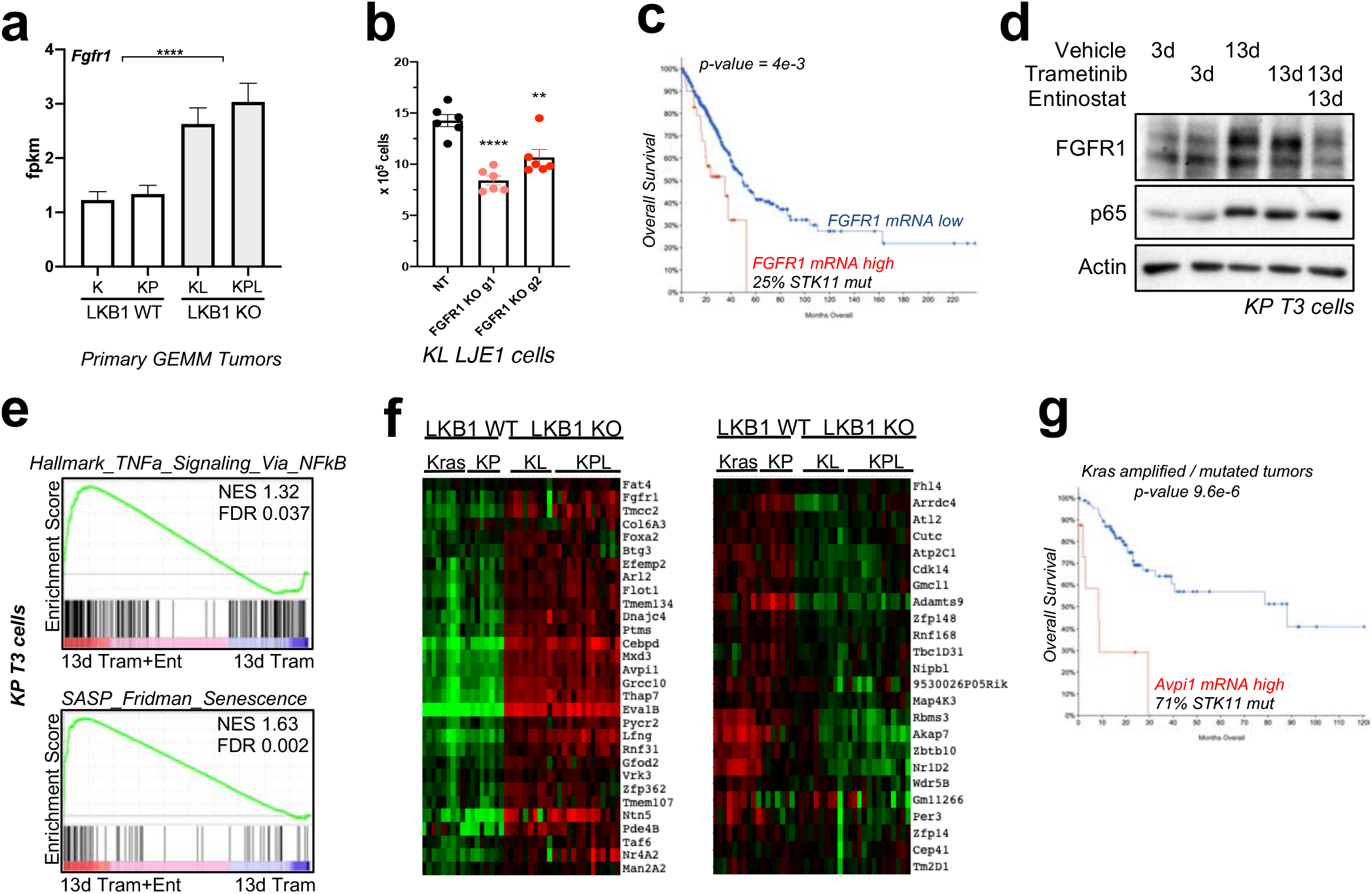
HDAC3 and NKX2-1 co-regulate a set of target genes which correlate with LKB1 mutation status. (A) *Fgfr1* mRNA expression across Kras (K) (n=9), KP (n=8), KL (n=9), and KPL (n=15) primary tumor RNA-seq data. (B) Proliferation assessment 5 days after plating NT or FGFR1 KO KL LJE1 cells (n=6). (C) Overall survival data from all patients in the Firehose Legacy LUAD TCGA dataset separated between *Fgfr1* mRNA high versus low tumors. (D) Western blot analysis of FGFR1 and p65 in protein lysates from KP T3 cells treated with Vehicle, 10nM Trametinib, or 1uM Entinostat for 3 or 13 days. (E) GSEA plots of the “Hallmark TNFa Signaling Via NFkB” and “SASP Fridman Senescence” gene sets queried against RNA-seq data comparing 13 day Trametinib+Entinostat (Tram+Ent) to 13 day Trametinib (Tram) conditions in KP T3 cells. (F) Heatmaps of RNA-seq data showing FPKM read counts from Kras (n=9), KP (n=8), KL (n=9), and KPL (n=15) primary tumors for the 43 TIER genes which are HDAC3 ChIP-seq target genes, and are expressed in an LKB1-dependent manner (genes in red boxes in Figure 4E). (G) Overall survival data from patients with Kras amplified or mutant tumors comparing tumors with or without high *AVPI1* mRNA in the Firehose Legacy LUAD TCGA dataset. Values are expressed as mean ± s.e.m. ** p-value < 0.01, **** p-value < 0.0001 determined by two-tailed student’s t-test.

## MATERIALS AND METHODS

### Cell culture and cell lines

All cell lines were incubated at 37 °C and were maintained in an atmosphere containing 5% CO_2_. 634T cell were a kind gift from Dr. Kwok Wong. KP T3 cells were published in Hollstein et al. *Cancer Discovery*, 2019. Cells were tested for Mycoplasma (Lonza) using manufacturer’s conditions and were deemed negative. Cells were grown in Dulbecco’s modified Eagles medium (DMEM) plus 10% fetal bovine serum (Gibco) and were continuously maintained under antibiotic selection for stable cell lines. Proliferation assays were performed by plating 2×10^3 cells per well of a 6-well plate, and cells were counted 5 days post-plating. Trametinib was used at 10nM, Entinostat was used at 1uM (long-term treatments) or 2uM (6h treatments), RGFP966 was used at 10uM, and TSA was used at 0.5uM. Treatments were 6h, 3d, or 13d. For long-term treatments, media was changed and fresh drug added every 2 days.

### Generating primary tumor cell lines

To generate KL LJE1 and KL LJE7 cell lines, individual primary tumors were dissected from the lungs of KL mice, mechanically dissociated, then digested for 45min in digestion media (10% FBS, pen/strep, 1mg/mLCollagenase/Dispase (Roche) in DMEM) at 37 °C. Cells were strained through 70uM nylon cell strainer, spun at 2000rpm 5min, resuspended in 1mL complete media plus 5uL Fungizone (Lifetech) and plated in a 24 well dish. 24h later, cells were infected by adding 1mL T-antigen-expressing lentivirus to each well. 24h later, viral media was removed and replaced with complete media with Fungizone. Cells were cultured in Fungizone for 4 weeks, then Epcam+ sorted.

### CRISPR/Cas9 studies

Small Guide RNAs (sgRNAs) targeting mouse HDAC3 were selected using the optimized CRISPR design tool (http://crispr.mit.edu). The gSR gRNA sequence targeting NKX2-1 was obtained from Sanchez-Rivera et al. *Nature* 2014 (46), and the other gRNA targeting NKX2-1, g2, was designed with the GPP sgRNA designer (https://portals.broadinstitute.org/gpp/public/analysis-tools/sgrna-design). gRNAs targeting FGFR1 and p65 were designed with the Benchling program (https://www.benchling.com/crispr/). Guides with high targeting scores and low probability of off-target effects were chosen. At least three independent sgRNA sequences were tested for each gene. Oligonucleotides for sgRNAs were synthesized by IDT, annealed in vitro and subcloned into BsmbI-digested lentiCRISPRv.2-puro (Addgene 52961) or lentiCRISPRv.2-hygro (Addgene 98291). Validation of guide specificity was assessed by Western blot of low-passage cells. Oligonucleotide sequences are listed in Supplemental Table S1.

### Transfection assays

Cells were transfected overnight in 10cm plates with N-GFP-RelA (Addgene #23255) or C-Flag-Rela (Addgene #20012) using 25μL Lipofectamine and 4.5μg plasmid DNA in 600μL Optimem total. The next day, media was changed. Two days after transfection, cells were treated with Vehicle (DMSO) or 2μM Entinostat for 6 hours and then collected.

### Lentiviral production and Titering

Lentiviruses made from pLentiCRISPRv.2 were produced by co-transfection of the lentiviral backbone constructs and packaging plasmids pSPAX2 (Addgene 12260) and pMD2.G (Addgene 12259). Lipofectamine 2000 (Thermo Fisher Scientific) was used as a transfection reagent at a ratio of 3:1 lipofectamine/DNA. Viral supernatant was collected from 293 cells 48 post-transfection, 0.45um-filtered, supplemented with polybrene, and applied to destination cells for 24h. Destination cells were allowed to recover from infection 24h before being subjected to selection with 2ug/ml puromycin or 150ug/mL hygromycin. Resulting stably transduced lines were frozen down immediately after selection. Large-scale viral preps of Lenti Pgk-Cre (a gift from Tyler Jacks) were made by the University of Iowa Viral Vector Core. Titering: Lentiviral preps for mouse experiments (PGK-Cre) were functionally titered by transduction of a reporter line (293-LSL-GFP), which turns on expression of GFP upon Cre-mediated recombination and allows quantitation of functional titers derived from the percent of GFP-positive cells.

### Mouse Studies

All procedures using animals were approved by the Salk Institute Institutional Animal Care and Use Committee (IACUC). All mice were maintained on the FVB/n background. **<u>Kras</u>** (*Kras*^LSLG12D/+;^ *R26* LSL;luc/luc); **<u>KL</u>** (*Kras* LSLG12D/+; *Lkb1* fl/fl;*R26* LSL;luc/luc), **<u>KP</u>** (*Kras* LSLG12D/+; *p53* fl/fl;*R26* LSL;luc/luc), and **<u>KPL</u>** (*Kras^LSLG12D/+^*^;^ *Lkb1^fl/fl^*;*p53^fl/fl^*;*R26^LSL;luc/luc^*) mice in FVB/n have been previously described(23) (21). The *Hdac3^fl/fl^* conditional floxed mouse has also been described (19). In this study, *Hdac3^fl/fl^* was crossed into the FVB/n K background before crossing into the KL or KP genotypes to generate KL-HDAC3^fl/fl^ and KP-HDAC3^fl/fl^ experimental mice. All experiments used a mixture of female and male mice. Lentivirus expressing Cre recombinase (4×10^5 pfu/mouse) was delivered by intratracheal intubation to each mouse to initiate lung tumorigenesis, according by the protocol of DuPage (47). Experimental endpoint was defined across experiments as the time point at which the experimental cohorts of KL or KP mice reached BLI tumor burden of 10^8 mean photon flux, or earlier as indicated. At endpoint, all mice in that experiment were collected at that point. All animals at experimental endpoint were included for analysis of lung tumor burden and tumor size analysis. No animals were excluded from longitudinal BLI measurements and graphs.

### BLI imaging

Bioluminescent imaging was performed biweekly using an IVIS Spectrum (Caliper Life Sciences) using Living Image software (Perkin Elmer). Mice were injected intraperitoneally with 150mg/kg D-luciferin (Caliper Life Science, Hopkinton MA), anesthetized with isoflurane and imaged both ventrally and dorsally 10 minutes post luciferin injection. The total lung photon flux for each animal is calculated by the combination of ventral and dorsal photon flux calculated within a region of interest (ROI) encompassing the thorax.

### *In vivo* HDAC inhibitor treatment

Mice were intratracheally intubated with Lentivirus expressing Cre recombinase to initiate tumorigenesis and imaged every two weeks starting 4 weeks post-Cre until average BLI was >5×10^7 mean photon flux. Mice were then randomized and treatment was initiated. Entinostat was diluted to 1 mg/mL in Vehicle (0.5% Methyl cellulose in water), vortexed 10 minutes, and administered daily at 10mg/kg by oral gavage at ~9am. RGFP was diluted to 0.8mg/mL in Vehicle (sequential addition of 7% Tween-20, 0.9% saline), vortexed 30 minutes, and administered daily at 10mg/kg by i.p. injection. On the 5^th^ day of treatment, endpoint material was collected 2-3 hours post-drug administration.

### Immunohistochemistry and image analysis

Lungs from mice were collected at each experimental endpoint as noted in the Fig.s, and were fixed in formalin for 18-22hrs, transferred to 70% ethanol and paraffin-embedded (FFPE) at the Tissue Technology Shared Resources at UCSD. 5μm sections from FFPE tissues were prepared and stained with hematoxylin and eosin. For immunohistochemistry, slides were deparaffinized and rehydrated, and antigen retrieval was performed for 13min at high heat (~95**°** C) in citrate buffer. Endogenous peroxidase activity was quenched with 10min hydrogen peroxide in methanol. Using the ImmPress HRP Ig (Peroxidase) Polymer Detection Kits (Vector Labs), slides were blocked, incubated overnight with primary antibody diluted in blocking buffer, and secondary antibody steps were carried out according to the manufacturer’s instructions. Staining was visualized with ImmPACT DAB peroxidase substrate (Vector Labs, SK-4105), and further counterstained with hematoxylin, dehydrated through ethanol and xylenes, and mounted with Cytoseal 60 (Thermo Scientific). H&E- and immunostained slides were scanned using a Perkin Elmer Slide Scanner (Panoramic MIDI Digital Side Scanner) for further downstream analysis using the Panoramic Viewer software and Inform v2.1 image analysis software (Cambridge Research and Instrumentation).

### Lung tumor burden

Total lung tumor burden was quantitated from H&E sections using Inform v2.1 image analysis software (Cambridge Research and Instrumentation) in a non-biased manner. In brief, the Trainable Tissue Segmentation method was trained to identify tumor, normal lung, vessel and space. This program was then applied to all H&E images, and each of the resulting mapped images was then screened to verify that accurate tissue segmentation had occurred. The quantitation data from this analysis was then used to calculate the percentage of tumor area as normalized to total lung area (tumor area + normal lung area).

### Tumor Size Quantitation

Quantitation of each individual tumor was measured from H&E sections using morphometric analysis in Panoramic viewer software (Perkin Elmer), which calculates the size of each identified tumor by area in squared microns. The area of all tumors found in the 5 lobes of each mouse was exported and compiled to plot the number of tumors per mouse, and the average size of every tumor in the cohort.

### mRNA preparation and qRT-PCR

mRNA was collected from cells harvested within 2 passages post-thaw. mRNA was prepared using the Quick-RNA Miniprep kit (Zymo Research), including DNase treatment. cDNA was synthesized from 2 μg of RNA using SuperScript III (Life Technologies), and qPCR was carried out with diluted cDNA, appropriate primers, and SYBR Green PCR master mix (ThermoFisher Scientific) using a C1000 Thermal Cycler (BioRad). Relative mRNA levels were calculated using the 2− Ct method, using *Tbp* as an internal control.

### mRNA-sequencing

RNA was isolated using the Quick-RNA MiniPrep kit (Zymo Research), including a DNase treatment. RNA integrity (RIN) numbers were determined using the Agilent TapeStation prior to library preparation. mRNA-seq libraries were prepared using the TruSeq RNA library preparation kit (version 2), according to the manufacturer’s instructions (Illumina). Libraries were quantified, pooled, and sequenced by single-end 50 base pairs using the Illumina HiSeq 2500 platform at the Salk Next-Generation Sequencing Core. Raw sequencing data were demultiplexed and converted into FASTQ files using CASAVA (version 1.8.2).

### Bioinformatic analysis of RNA-seq data

Sequenced reads were quality-tested using the online FASTQC tool (http://www.bioinformatics.babraham.ac.uk/projects/fastqc) and aligned to the mouse mm10 genome using the STAR aligner version 2.4.0k (48). Raw gene expression was quantified across all annotated exons using HOMER (49), and differential gene expression was carried out using the getDiffExpression.pl command. Differentially expressed genes were defined as having a false discovery rate (FDR) <0.05 and a log2 fold change >0.5.

GSEA was carried out with the GenePattern interface (https://genepattern.broadinstitute.org) using preranked lists generated from FDR values. Queried datasets used were (1) GSEA Gene Set “Hallmark_TNFA_Signaling_Via NFKB” M5890, (2) SASP Fridman Senescence, and (3) gene lists from genes differentially expressed upon Tamoxifen-driven NKX2-1 KO in Kras tumors (37). Heatmaps were generated by clustering using the Cluster 3.0 program (log2 transform data, center genes, Hierarchical clustering with average linkage) (50), and then visualized with Java TreeView version 1.1.6r4 (51).

### ChIP-sequencing

#### Primary tumors

Individually dissected, flash frozen primary tumors were combined from 3 different mice into one pool of 130mg of primary tumors per replicate per genotype. Equivalent masses of tumors were used from each of the three mice to ensure equal representation. Two independent pools of tumors per genotype were processed separately to generate two biological replicate pool of crosslinked, sonicated chromatin for ChIP. 4 independent ChIPs were performed on each pool of sonicated chromatin, and then pooled together to generate one replicate for ChIP-sequencing. To crosslink, tumors were dounce homogenized in crosslinking buffer (1% Formladehyde in PBS) and incubated with end-over-end rotation for 15min at room temperature, and then quenched with 2.5M glycine 5min. Samples were spun at 600g for 5min, washed with cold PBS, and resuspended in ChIP buffer (RIPA) (see “Immunoprecipitation” for recipe) with protease inhibitors. Samples were sonicated in a Covaris LE 220 for 8min (Duty Factor 2, 105 Watts, 200 cycles/burst), spun down, and the supernatant saved. For each ChIP, 100uL lysate was combined with 900uL ChIP buffer, while 50uL was used for Input. 10ug of Hdac3 ab7030 antibody and 2ug H3K27ac ab4729 antibody was used for each ChIP. Lysate was incubated overnight with antibody. 20uL washed and pre-blocked Protein A Dynabeads were incubated 2hrs rotating with each sample at 4degC. Washes were performed with 5min incubations of each buffer while rotating at 4degC. Samples were washed 3X with cold ChIP buffer, 1X with room temperature ChIP buffer, and 1X with room temperature TE pH 8, and then spun down. Elution of ChIP and Input samples was done by incubating samples with Elution buffer (50mM Tris/Hcl pH 7.5, 10mM EDTA, 1% SDS) overnight at 65degC. Beads were pelleted and discarded, and 200uL of eluate was combined with 194uL low-EDTA TE and 100ug proteinase K, and incubated 2hrs at 37degC. 8uL RNase A was added and samples incubated 30min at 37degC. Minelute PCR purification kit (Qiagen 28006) was used to isolate DNA, which was eluted in 15uL EB at 55degC. 4 ChIPs were combined into one sample for ChIP-sequencing.

#### KL LJE1 cells

ChIP-seq was carried out on DSG+Formaldehyde crosslinked, sonicated nuclear extracts. Cells were washed in PBS and then crosslinked by 30min incubation in 2mM DSG (Di(N-succinimidyl) glutarate, Thermo Fisher NC0054325). Aspirate, and incubate 15min with 1% formaldehyde, before 5min of quench with 125mM glycine. Cells were washed in cold PBS, scraped, and spun down, and washed again in PBS before nuclei isolation. Nuclei were isolated by resuspension in CiA NP-Rinse 1 (50mM HEPES, 140mM NaCl, 1mM EDTA, 10% glycerol, 0.5% NP-40, 0.25% Triton X100), incubated 10min at 4degC with end-over-end rotation, then centrifuged at 1,200g for 5min at 4degC. Samples were then resuspended in CiA NP-Rinse 2 (10mM Tris pH 8.0, 1mM EDTA, 0.5mM EGTA, 200mM NaCl), incubated 10min at 4degC with end-over-end rotation, and centrifuged at 1,200g for 5min at 4degC. Tubes were washed 2X with Covaris Shearing Buffer (0.1% SDS, 1mM EDTA pH 8, 10mM Tris HCl pH 8) to remove salt, centrifuged at 1,200g at 4degC 3min. Samples were diluted to a concentration of 2.5×10^6^ cells/130uL in ChIP buffer (RIPA) (50mM Tris-HCl pH 7.5, 140mM NaCl, 1mM EDTA, 1% Triton X-100, 0.1% NaDOC (sodium deoxycholate), 0.1% SDS) with protease inhibitors, and sonicated in a Covaris LE 220 for 8min (Duty Factor 2, 105 Watts, 200 cycles/burst). Sonicated material was spun down and supernatant was used for ChIP. Lysate from 5 million cells was diluted in ChIP buffer to 1mL final volume. 50uL was used for Input. 10ug of Hdac3 CST-85057 antibody and 2ug H3K27ac ab4729 antibody was used for each ChIP. Lysate was incubated overnight with antibody. 20uL washed and pre-blocked Protein A Dynabeads were incubated 2hrs rotating with each sample at 4degC. Washes were performed with 5min incubations of each buffer while rotating at 4degC. Samples were washed 3X with cold ChIP buffer, 1X with room temperature ChIP buffer, and 1X with room temperature TE pH 8, and then spun down. Elution of ChIP and Input samples was done by incubating samples with Elution buffer (50mM Tris/Hcl pH 7.5, 10mM EDTA, 1% SDS) overnight at 65degC. Beads were pelleted and discarded, and 200uL of eluate was combined with 194uL low-EDTA TE and 100ug proteinase K, and incubated 2hrs at 37degC. 8uL RNase A was added and samples incubated 30min at 37degC. Minelute PCR purification kit (Qiagen 28006) was used to isolate DNA, which was eluted in 15uL EB at 55degC.

### Bioinformatic analysis of ChIP-seq data

Sequenced reads were aligned to the mouse mm10 genome using the STAR aligner version 2.4.0k (48). HOMER (49) was used for data processing. For KL LJE1 cell line ChIP-seq data, peaks were called using the getDifferentialPeaksReplicates.pl command using HDAC3 ChIP-seq data from NT cells as target (−t), HDAC3 ChIP-seq data from HDAC3 KO cells as background (−b), and Input sequencing data from NT cells as input (−i), with -style factor and −F 3. For primary tumor ChIP-seq data, peaks were called for each replicate individually using the findPeaks command with parameters-style factor −F 3, using HDAC3 ChIP-seq as target and Input sequencing data as Input (−i). Peaks were merged using the mergePeaks command to generate a consolidated file containing all HDAC3 ChIP-seq peaks identified in KL and KP tumors. The getDifferentialPeaks command with −F 3 -same was used to identify peaks bound in both KL and KP tumors. The annotatePeaks.pl command with the -ghist -hist 25 option was used to visualize binding at each peak independently across samples, and Java TreeView was used to visualize the output. The annotatePeaks.pl command with -hist 25 was used to plot average reads across all peaks relative to peak center for each replicate separately. BedGraph files were also generated and visualized with Integrative Genomics Viewer (IGV) version 2.5.1.

### Homer motif enrichment analysis

Homer motif enrichment analysis: http://homer.ucsd.edu/homer/motif/.

### Web-based analysis tools

Pathway analysis was performed with Enrichr: http://amp.pharm.mssm.edu/Enrichr. 4-way Venn diagrams were plotted using Venny 2.1 (Oliveros, J.C. (2007–2015) Venny. An interactive tool for comparing lists with Venn’s diagrams, (https://bioinfogp.cnb.csic.es/tools/venny/). Area-proportional Venn diagrams were plotted using BioVenn (52) (https://biovenn.nl).

### Western blots

Protein lysates in CST buffer (20mM Tris pH 7.5, 150mM NaCl, 1% Triton X-100, 50mM NaF, 1mM EDTA, 1mM EGTA, 2.5mM PyroPhosphate, 2mM beta-glycerol-phophate, 1mM orthovanadate, 0.01mM Calyculan A) with protease inhibitors were equilibrated for protein levels using a BCA protein assay kit (Pierce), resolved on 8% SDS-PAGE gels, and transferred to membrane. Membranes were blocked in milk, incubated overnight at 4degC in diluted primary antibody, washed with TBS-T, incubated 1hr in secondary antibody diluted in in TBS-T plus milk, washed in TBS-T, and developed using SuperSignal ECL. Secondary antibodies used were anti-rabbit (Millipore AP132P) and anti-mouse (Millipore AP124P). Nuclear fractions were isolated using a NE-PER nuclear and cytoplasmic extraction kit (Thermofisher) under manufacturers conditions. Quantitation was performed with ImageJ (Rasband, W.S., ImageJ, U. S. National Institutes of Health, Bethesda, Maryland, USA, https://imagej.nih.gov/ij/, 1997-2018).

### Immunoprecipitation

Immunoprecipitation was carried out on DSP-crosslinked, sonicated nuclear lysates. Cells were washed in PBS and then crosslinked by 30min incubation in 1mM DSP (dithiobis(succinimidyl propionate), Thermo Scientific 22585), followed by 5min of quench with 2.5M glycine. Cells were washed in PBS, scraped, and spun down, and washed again in PBS before nuclei isolation. Nuclei were isolated by resuspension in CiA NP-Rinse 1 (50mM HEPES, 140mM NaCl, 1mM EDTA, 10% glycerol, 0.5% NP-40, 0.25% Triton X100), incubated 10min at 4degC with end-over-end rotation, then centrifuged at 1,200g for 5min at 4degC. Samples were then resuspended in CiA NP-Rinse 2 (10mM Tris pH 8.0, 1mM EDTA, 0.5mM EGTA, 200mM NaCl), incubated 10min at 4degC with end-over-end rotation, and centrifuged at 1,200g for 5min at 4degC. Tubes were washed twice with Covaris Shearing Buffer (0.1% SDS, 1mM EDTA pH 8, 10mM Tris HCl pH 8) to remove salt, centrifuged at 1,200g at 4degC 3min. Samples were diluted to a concentration of 2.5×10^6^ cells/130uL in ChIP buffer (RIPA) (50mM Tris-HCl pH 7.5, 140mM NaCl, 1mM EDTA, 1% Triton X-100, 0.1% NaDOC (sodium deoxycholate), 0.1% SDS) with protease inhibitors, and sonicated in a Covaris LE 220 for 8min (Duty Factor 2, 105 Watts, 200 cycles/burst). Sonicated material was spun down and supernatant was used for IP: 400uL material was incubated with 3uL NKX2-1 antibody (Abcam ab76013) per IP overnight with rotation at 4degC. 20uL prewashed Protein A Dynabeads were added per tube, and incubated 4hrs at 4deg with rotation. Samples were washed 5x with CST buffer (see western blot section) before adding 25uL 6x loading dye and 50uL CST per tube, and eluting by boiling for 5min. Input and IP samples were subsequently assessed by western blot.

### TCGA analysis of Firehose LUAD dataset

The results shown are in whole based upon data generated by the TCGA Research Network: http://cancergenome.nih.gov/. TCGA datasets were queried using cBioPortal (www.cbioportal.org) (53) (54). Methods for data generation, normalization and bioinformatics analyses were previously described in the TCGA LUAD publication (Cancer Genome Atlas Research 2014). mRNA data used for this analysis was RNA Seq V2 RSEM with *z*-score thresholds of 1.8.

### Antibodies and reagents

#### Western blotting

Antibodies from Cell Signaling Technologies (Denvers, MA USA) were diluted 1:1,000: Hdac3 CST-85057, p65 CST-6956, acetyl-NF-kB p65 (Lys310) CST-12629, Hdac4 CST-15164, Fgfr1 CST-9740. Nkx2-1 was from Abcam ab76013 and was diluted 1:1,500. From Sigma-Aldrich, anti-actin (#A5441) was diluted 1:10,000.

#### Immunohistochemistry

Abcam antibodies ab5690 was used at 1:150 to detect CD3, ab198505 was used at 1:100 to detect Cxcl5/6. Cell Signaling Technologies CST-70076 was used at 1:250 to detect F4/80. R&D Systems AF2225 was used at 5ug/mL to detect NKp46/NCR1. BioXcell BE0075-1 was used at 1:1,000 to detect Ly6g. SantaCruz sc-6246 was used at 1:50 to detect p21.

#### ChIP

Hdac3 from Abcam ab7030 was used on primary tumors, and Hdac3 from Cell Signaling Technologies CST-85057 was used on KL LJE1 cells.

#### IP

Nkx2-1 raised in rabbit from (Abcam ab76013) was used to immunoprecipitate, and HDAC3 raised in mouse (CST-3949) was used to detect co-immunoprecipitated HDAC3.

### Statistical analyses

Statistical analyses are described in each Fig. and were all performed using Graph Pad Prism 9. Results are expressed as mean ± s.e.m. unless otherwise indicated.

## Acknowledgements

This study was supported by grants to R.J.S. from the NIH (R35CA220538, P01CA120964) and The Leona M. and Harry B. Helmsley Charitable Trust grant #2012-PG-MED002. L.J.E. was supported by a postdoctoral fellowship from the American Cancer Society (PF-15-037-01-DMC). S.N.B was supported by NCI training grant 5T32CA009370 to the Salk Institute Cancer Center and NCI 5F32CA206400. This work was supported by the NGS and the Razavi Newman Integrative Genomics and Bioinformatics Core Facilities of the Salk Institute with funding from the NIH-NCI CCSG: P30 014195, the Chapman Foundation, and the Helmsley Charitable Trust. Tissue Technology Shared Resource is supported by a National Cancer Institute Cancer Center Support Grant (CCSG Grant P30CA23100). We thank Dr. Scott Hiebert (VUMC) for the gift of the Hdac3 floxed mice and Dr. Diana Hargreaves and Dr. Jarrett Remsberg for support with ChIP-seq assay development.

## Author contributions

L.J.E designed all experiments, performed all experiments except as noted here, and wrote the paper. S.D.C., S.N.B, and J.T.B. assisted with cloning and KO cell line generation and validation. S.N.B. and J.T.B. assisted with *in vivo* drug dosing studies. S.D.C. assisted with qRT-PCR. D.S.R. assisted with animal husbandry. T.J.R. assisted with genotyping and BLI imaging. L.J.E. and R.J.S. conceived the project, and R.J.S. supervised the project and wrote and edited the paper.

## Data availability

RNA-seq and ChIP-seq data have been deposited to the Gene Expression Omnibus (GEO) data repository with accession number GSE164759. Previously published sequencing data that were reanalyzed here is available under the GEO accession code GSE36473. Source Data are provided for all experiments. Other data that support the findings of this study are available on request from the corresponding author upon reasonable request.

